# Prevalence of antimicrobial resistance genes and integrons in commensal Gram-negative bacteria in a college community

**DOI:** 10.1101/683524

**Authors:** Julia Rubin, Kaitlyn Mussio, Yuqi Xu, Joy Suh, Lee W. Riley

## Abstract

Although the human intestinal microbiome has been shown to harbor antimicrobial drug-resistance genes (ARG), the prevalence of such genes in a healthy population and their impact on extraintestinal infections that occur in that community are not well established. This study sought to identify ARG prevalence and their mobile elements in the intestines of a healthy community population at a California university, and compared these genes to those found in uropathogenic *Escherichia coli* isolated from patients with community-acquired urinary tract infection (CA-UTI). We isolated Gram-negative bacteria (GNB) from fecal samples of healthy volunteers and screened them by polymerase chain reaction (PCR) for ARG encoding resistance against ampicillin (AMP), trimethoprim-sulfamethoxazole (TMP-SMX), gentamicin (GENT), and colistin (COL). We found antimicrobial resistant GNB from 85 (83%) of 102 non-redundant rectal swab samples. Sixty-seven (66%) of these samples contained ß-lactamase genes (*bla*_TEM_, *bla*_SHV_, *bla*_CTX-M_, *bla*_OXA,_ *bla*_OXY_), dihydrofolate reductase (DHFR) genes (*dhfr-A17*, *dhfr-A7*, *dhfr-A5*, *dhfr-A21*, *dhfr-A1*, *dhfr-A15*, and *dhfr-B3*), and aminoglycoside resistance genes (*aadA5*, *aadA1*, and *aadB*). Integron sequences were found in 37 fecal samples. These genes were found in 11 different GNB species isolated from the fecal samples. The same ARG were found in *E. coli* strains isolated from patients with CA-UTI examined at the college outpatient health clinic. The high prevalence of clinically-common ARG and integrons harbored by GNB in the intestine of a healthy population suggest that human intestines may serve as a major reservoir of these mobile ARG that appear in *E. coli* strains causing extraintestinal infections in the same community.

**Importance:** Increasing frequency of antimicrobial resistance (AMR) in human pathogenic bacteria has compromised our ability to treat infections. Since mobile antibiotic resistance genes (ARG) are readily exchanged between different species of bacteria through horizontal gene transfer, there is interest in investigating sources of these genes. The normal intestinal flora has been shown to contain a wide variety of ARG, which may have been introduced via food-containing AMR bacteria. We sought to assess the prevalence of ARG carriage in the intestines of a healthy population and determine if these ARG are found in *E. coli* strains that cause community-acquired urinary tract infection (CA-UTI) in the same community. Our findings indicate that the human intestine may serve as an important reservoir as well as a site in which ARG are transferred into *E. coli* that cause UTI. Further research is needed to reduce ARG carriage and devise new strategies to prevent AMR infections.

## Introduction

The emergence of antibiotic resistant bacteria and antibiotic resistance genes (ARG) has become a growing public-health concern worldwide. Gram-negative bacteria (GNB) including *Enterobacteriaceae* and glucose non-fermenter species are implicated in a large number of community and healthcare-associated infections. They are also common residents of the human intestinal microbiota as well as of the environment (1–10). Human and veterinary medicines rely on the use of antibiotics to treat infections caused by bacterial pathogens. Antibiotics are also used in food animal production as growth promoters. Extensive use of antibiotics over the last several decades has led to the rapid emergence and spread of resistant microbes, especially among enteric bacteria (11, 12). Due to the nature and complexity of the microbial communities harbored in our intestinal environment, the human gut has been recognized as a potential reservoir of ARG (3, 13, 14). However, ARG are also found in food animal intestines as well as in a variety of produce items humans eat (12, 13, 15–17, 21–34). Thus, food may serve as sources of ARGs that enter the human intestine. The prevalence of ARG in healthy human intestine and their impact on human infections is not well characterized.

The ability of bacteria to disseminate mobile ARG via horizontal gene transfer (HGT) has resulted in the rapid acquisition of resistance by enteric bacterial pathogens. ARG mobile elements include integrons, transposons, and plasmids. Integrons, which permit the simultaneous integration of multiple exogenous gene cassettes, represent an important vehicle for the rapid horizontal transfer of resistance across bacterial populations and thus could contribute to the sudden increase in prevalence of multidrug-resistant infections in a community (19, 20). In particular, class 1 integrons are the most ubiquitous class of integrons in enteric bacteria and have been found in all common pathogens including *Escherichia*, *Klebsiella*, *Salmonella*, *Shigella*, and other disease-causing Enterobacteriaceae. More than 70 different gene cassettes conferring resistance to most of the known β-lactam drugs, aminoglycosides, trimethoprim, rifampicin, chloramphenicol, and erythromycin have been reported in class I integrons (11). This is of further concern due to the increasing prevalence of extended spectrum beta lactamase (ESBL)-producing GNB, which are difficult to treat and often resistant to other families of antibiotics, in particular trimethoprim-sulfamethoxazole and fluoroquinolones (7). Furthermore, the recent global spread of GNB containing plasmid-mediated colistin resistance has greatly challenged clinical management of infections caused by them (21). For example, colistin remains an important therapeutic agent for infections caused by carbapenem-resistant Enterobacteriacaea (CRE), in which there are limited alternative treatment options (22).

Previous studies have shown food and the environment to contain ARG found in pathogenic bacteria (12, 23, 24). Environmental reservoirs of ARG-containing pathogens include lakes and rivers (18), wastewater treatment plants (25, 26), houseflies (27), livestock (12, 28, 29), soil and manure (29, 30), retail meat products (12, 31–33), companion animals (5, 6, 8, 28, 34), alfalfa sprouts (35), retail spinach (15), and other vegetables (36). The findings from these studies indicate that human health may be impacted by uptake of pathogens from the environment, many of which harbor ARG on mobile genetic elements. Thus, humans may be acquiring pathogens as well as transmissible ARG through food and other environmental hosts.

A recent study by Yamaji et. al compared ARG and genotypes of *E. coli* isolates from patients with urinary tract infection (UTI) in a California university community to those of *E. coli* isolated from meat (pork, chicken, beef, and turkey) obtained from retail stores in the surrounding area (31). They found that despite 12 shared genotypes (sequence types) between humans and retail meat *E.coli* isolates, human isolates contained more ARG than did meat isolates (31). These findings led us to speculate that people may acquire uropathogenic *E. coli* from external sources (food or environment) but that these *E. coli* strains then may acquire ARG from commensal bacteria already in the gut. Therefore, we sought to understand the diversity and abundance of mobile ARG in the gut of healthy humans, and to elucidate potential links between ARG harbored in commensal gut bacteria and the acquisition of those ARG by pathogenic GNB.

## Materials and methods

### Sample collection

We prospectively cultured fecal samples from 113 healthy volunteers at a university campus in northern California between June and October 2017. Eligible participants included those between 18 and 65 years of age, with no medical history of urinary tract corrective surgery or abnormality, or bladder catheterization or hospitalization within the 6 months prior to sample collection.

At recruitment, participants were provided a pre-addressed collection kit containing a Cary-Blair transport media rectal swab (Becton Dickinson BBL™), two biosafety bags, and detailed collection instructions. Each kit also included a questionnaire regarding antibiotic use, history of UTI, and diet and lifestyle characteristics. Participants were instructed to send the rectal swab back to the laboratory via USPS mail immediately after collection. Once delivered, the study coordinator analyzed samples within 48 hours.

### Fecal sample analysis

Fecal swab tips were placed in a 1.5-ml Eppendorf tube containing 1mL of Luria-Bertani broth and vortexed for 60 seconds. A 10μl aliquot of fresh fecal material was dilution streaked onto MacConkey agar plates and incubated overnight at 37°C. All samples were screened for resistance against four antimicrobial agents: ampicillin (AMP) (32μg/ml), gentamicin (GENT) (16μg/ml), trimethoprim-sulfamethozaxole (TMP-SMX) (4-76 μg/ml), and colistin (COL) (2μg/ml), as well as on one MacConkey agar plate containing no drug. All antimicrobial agents were dissolved in nuclease free water and filter sterilized. Dimethyl sulfoxide (DMSO) was used to prepare TMP-SMX solution (final concentration <5% DMSO). Dihydrofolate reductase (DHFR) mediates trimethoprim resistance. We tested trimethoprim-sulfamethozale in this study as this combination drug is commonly used to treat UTI and other bacterial infections. Interpretive criteria from the Clinical Laboratory Standard Institute (37) or from literature recommendations (38) were used to determine resistance. *E. coli* 25922 (ATCC) was used as a reference strain.

### DNA extraction and ARG identification

Five bacterial colonies were randomly selected from each plate. If multiple colony morphologies were noted, all colonies with unique morphologies were sampled. If less than five colonies were present, all colonies were selected for analysis. Single colonies were selected and inoculated into 3 ml tryptic soy broth and incubated in a shaking incubator for 20 hours at 37°C. Basic procedures for DNA extraction by a freeze-thaw method were performed as previously described (15, 19). The 2ml aliquots of the cultures were centrifuged, and the pellets were re-suspended in a test tube with 350μl of distilled water, boiled for 10 min in a water bath, and then cooled on ice for 2 min. The samples were centrifuged for 2 min at 13,000 rpm, and the supernatants were stored at 20°C before they were subjected to PCR tests.

Two microliters of the resulting supernatant were used as template DNA in 25μl of PCR mixture. Bacterial isolates that grew in the presence of AMP were examined for ß -lactamase gene families by multiplex PCR as described previously (39). These β-lactamase gene families included: TEM variants (*bla*_TEM-1_ and *bla*_TEM-2_), an SHV variant (*bla*_SHV-1_), CTX-M (all variants), OXA variants (*bla*_OXA-1_, *bla*_OXA-4_, and *bla*_OXA-30_), and KPC variants (*bla*_KPC-1_ to *bla*_KPC-5_). Samples that showed bands corresponding to CTX-M universal variants were further examined for CTX-M group variants: CTX-M group 1 (*bla*_CTX-M-1_, *bla*_CTX-M-3_, and *bla*_CTX-M-15_), CTX-M group 2 (*bla*_CTX-M-2_), CTX-M group 9 (*bla*_CTX-M-9_ and *bla*_CTX-M-14_), CTX-M group 8/25 (*bla*_CTX-M-8_, *bla*_CTX-M-25_, *bla*_CTX-_M-26, and *bla*_CTX-M-39_ to *bla*_CTX-M-41_). Bacteria that grew on plates containing GENT or TMP-SMX were examined for 5’ and 3’ conserved sequences flanking the class I integron gene cassettes. Thus, the entire cassette sequences harbored in the integron were analyzed. If present, these gene cassettes were sequenced to detect the presence of *aad* and *dhfr* gene types for GENT and TMP-SMX, respectively. Bacterial colonies isolated from colistin-containing plates were examined for *mcr-1* and *mcr-2* genes as previously described (40). All isolates were speciated by 16S ribosomal RNA sequencing. Species diversity was calculated by the Shannon diversity index:

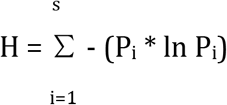

where:

H = the Shannon diversity index
P_i_ = fraction of the entire population made up of species i
S = numbers of species encountered
∑ = sum from species 1 to species S

Primers used in these procedures are noted in Table S1. PCR products were visualized on a 1.5% agarose gel stained with SYBRsafe DNA gel stain (Invitrogen) and visualized under UV transillumination.

### Sequencing Analysis

To identify gene or gene variants, we sequenced each PCR product that showed an electrophoretic band of an expected molecular weight. For amplicons over 1000 base pairs in length, bidirectional sequencing was performed. Each sequence was then compared against sequences in GenBank by BLAST (National Center for Biotechnology Information). Species were determined by the criteria of >98% sequence identity and genus by >95% sequence identity.

## Results

Between June 2017 and February 2018, we obtained 113 unique rectal swab samples. Of these, 11 exhibited no growth on a MacConkey control plate, and were not further analyzed. Of the 102 remaining samples, 76 (75%) contained GNB that exhibited resistance to AMP, 14 (14%) to GENT, 47 (46%) to TMP-SMX, and 10 (10%) to COL. Eighteen (18%) samples contained isolates that were sensitive to all drugs tested (Figure 1).

**Figure 1:**
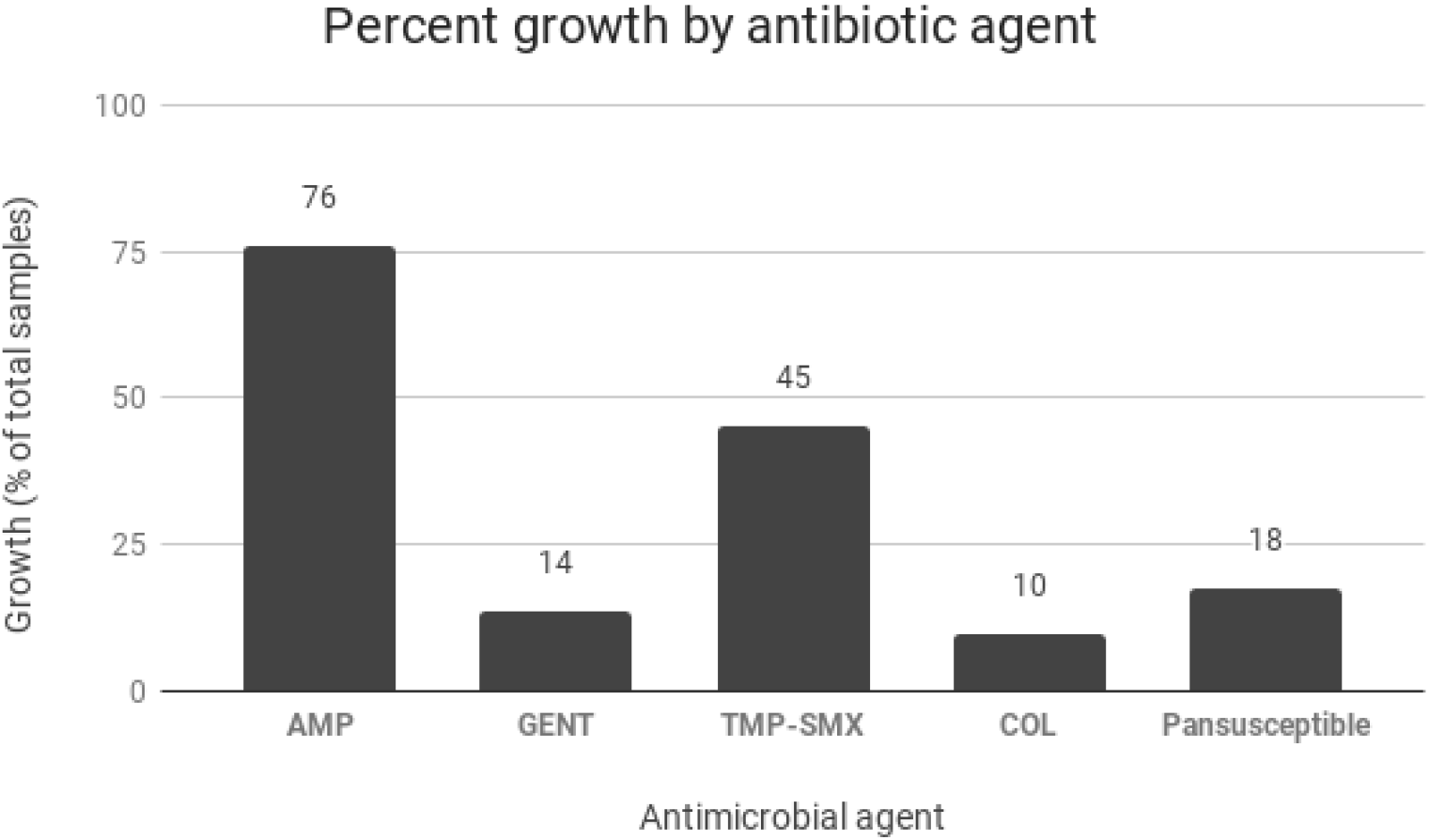
Bacterial growth (%) on MacConkey plates containing antimicrobial agents. Pansusceptible isolates were defined as those that exhibited growth on MacConkey control plate, but no growth in the presence of each of four antimicrobial agents tested.

Bacterial growth on at least one drug-containing plate was observed in 85 (83%) of 102 samples. Resistance to only 1 antimicrobial agent was observed in 33 (32%) of samples; 39 (38%) were resistant to two antimicrobial agents, and 13 (13%) were resistant to 3 or more agents (Table 1).

**Table 1:** Colony growth was observed in 82 fecal samples on MacConkey plates containing distinct types of antimicrobial agents. The number of types of antimicrobial agents to which a sample contained bacterial isolates that displayed resistance. AMP=ampicillin-resistant, TMP/SMX= trimethoprim-sulfamethoxazole-resistant, COL=colistin-resistant, GENT=gentamicin-resistant.

| Resistance to (number of) antimicrobial agents |  |
| --- | --- |
| Number, name of Antimicrobial agents | Number (%) of samples exhibiting phenotypic resistance |
| **0** | 19 (19%) |
| **1** | 31 (30%) |
| AMP | 27 (26%) |
| TMP/SMX | 2 (2%) |
| COL | 2 (2%) |
| **2** | 39 (38%) |
| AMP, TMP/SMX | 32 (31%) |
| AMP, COL | 3 (3%) |
| GENT, TMP/SMX | 1 (1%) |
| AMP, GENT | 3 (3%) |
| **3 or more** | 13 (13%) |
| AMP, GENT, TMP/SMX | 8 (8%) |
| AMP, TMP/SMX, COL | 4 (4%) |
| AMP, GENT, TMP/SMX, COL | 1 (1%) |
| **TOTAL** | 102 (100%) |

### Detection of ß-lactamase genes

Three hundred sixty-seven colonies obtained from 76 fecal samples containing AMP-resistant isolates were analyzed for the presence of β-lactamase genes. Of the 76 fecal samples, 68 (89%) contained at least one β-lactamase gene. Among the 102 total fecal samples, TEM variants (*bla*_TEM-1_, *bla*_TEM-116_) were detected in 57 (55%); 19 (18%) contained a SHV variant, 1 (1%) contained an OXA variant (*bla*_OXA-1_), and 15 (15%) contained at least one CTX-M variant. Upon further analysis of CTX-M subgroups, 6 (6%) had CTX-M group 1 (i.e. *bla*_CTX-M-15_), 2 (2%) had CTX-M group 2, and 4 (4%) contained CTX-M group 9 variants (i.e. *bla*_CTX-M-14_, *bla*_CTX-M-27_). Two (2%) contained more than one CTX-M group variant. Six samples contained a CTX-M gene that did not belong to any of the four CTX-M family members tested. Sequence analysis of these PCR products detected an OXY variant sequence in 4 of these samples (41, 42). Twenty-five isolates from 9 fecal samples contained confirmed CTX-M variants. No KPC or CTX-M group 8/25 gene types were detected in this study (Table S2).

### Detection of dihydrofolate reductase genes contained in class 1 integron gene cassettes

Two hundred twenty TMP-SMX-resistant colonies were recovered from 45 fecal samples; they were analyzed for the presence of a class 1 integron gene cassette to detect *dhfr* gene types. Gene cassettes were detected in 30 (67%) of 45 samples and ranged from 1-2.2 kilobases in size. We detected *dhfr* genes in 21 samples, consisting of the following gene variants: *dhfr-A17*, *dhfr-A7*, *dhfr-A5*, *dhfr-A21*, *dhfr-A1*, *dhfr-A15*, and *dhfr-B3*. *Dhfr-A17* was the most prevalent gene of this type, present in 10 (22%) of 45 samples (Table 3). Nine (30%) of the 30 samples that contained gene cassettes from phenotypically TMP-SMX resistant isolates did not contain any *dhfr* genes (Table S3).

### Detection of aminoglycoside adenyltransferase genes contained in class 1 integron gene cassettes

One of the 14 samples that initially showed resistance to GENT was unable to be cultured in growth media and thus was not further analyzed. Fifty-nine GENT-resistant colonies were isolated from 13 samples; they were analyzed for class 1 integron gene cassettes. Seven (54%) of the 13 samples contained the gene cassette; 6 (46%) of these contained at least 1 *aad* gene type. The following *aad* genes were detected: *aad*A5, *aad*A1, and *aad*B. Of the samples containing *aad* genes, *aad*A5 was detected in 5 of the 6 samples, while 1 sample contained both *aad*A1 and *aad*B (Table S4).

### Detection of mobile colistin resistance genes

Thirty-eight COL-resistant colonies were isolated from 10 samples and analyzed for *mcr-1* and *mcr-2* genes. No *mcr-1* or *mcr-2* was detected by PCR in these samples (Table S4).

### Detection of integron cassette sequences

We used PCR to amplify cassette sequences integrated in class 1 integrons in GNB strains that were resistant to GENT and TMP-SMX. Of 58 strains, 37 (64%) had integron sequences. In addition to the *dhfr* sequences described above, sequences from TMP-SMX-resistant bacteria were annotated in the NCBI BLAST database as *bla*_*OXA (−1, −2, −10, −48)*_, *bla*_*DHA*_, *bla*_*KPC*_, *bla*_*CTX-M (−2, −3, −15)*_, *bla*_NDM_, *bla*_CMY_, *bla*_IMP_, *bla*_GES_, *bla*_VIM_, *aadA1, aad*A2*, aad*A5*, aad*A6, *aac*A4*, aac*A6*, aac*A7*, aac*A8*, sul1*, and *mcr-1* (Table 2). In addition to the *aad* sequences described above, integron gene cassettes sequenced from GENT-resistant bacteria were annotated in the NCBI BLAST database as *bla*_OXA-1,_ *bla*_CTX-M (−3, −15)_, *bla*_NDM (−5, −9),_ *bla_CMY,_ bla*_IMP,_ *dhfr-A17*, and a *mcr variant* (Table 2, S3, S4).

**Table 2:** Number of fecal samples containing bacterial colonies from each genus. Colonies were isolated from bacteria resistant to indicated antimicrobial agent. Colony growth from fecal culture may represent multiple species. Class 1 integron cassette sequences were analyzed only for TMP-SMZ and GENT-resistant isolates.

| Genus | AMP-resistant (# of samples) | β-lactamase gene | TMP-SMX-resistant (# of samples) | *dhfr* gene in integron | Other ARG in integron in TMP-SMX-resistant strains | Growth on GENT (# of samples) | *aad* gene in integron | Other ARG in integron in GENT-resistant strains |
| --- | --- | --- | --- | --- | --- | --- | --- | --- |
| Escherichia or Shigella sp. | 38 | *bla*<sub>TEM</sub>, *bla*<sub>CTX-M</sub>, *bla*<sub>SHV</sub>, *bla*<sub>OXA</sub> | 34 | *dhfr*-A7, *dhfr*-A17, *dhfr*-B3, *dhfr*-A5 | *bla*<sub>OXA</sub> (-1, -2, -10, -48), *bla*<sub>CTXM</sub> (-3, -15), *bla*<sub>NDM</sub>-5, *bla*<sub>CMY</sub>, *bla*<sub>GES</sub>-11, *bla*<sub>DHA</sub>-1, *bla*<sub>IMP</sub>, *bla*<sub>SIM</sub>, *bla*<sub>VIM</sub>, *aacA6*, *aacA4*, *aacA7*, *aacA8*, *aadA5*, *aadA6*, *sul1*, *mcr*-1 | 11 | *aadA5*, *aadB*, *aadA1*, *aadA5* | *bla*<sub>NDM</sub> (-5, 9), *bla*<sub>OXA</sub>-1, *bla*<sub>CTXM</sub> (-3, -15), *bla*<sub>IMP</sub>, *bla*<sub>CMY</sub>, *dhfr*-17, *mcr* |
| Klebsiella sp. | 18 | *bla*<sub>TEM</sub>, *bla*<sub>SHV</sub>, *bla*<sub>OXY</sub> | 4 | *dhfr*-A1 | *bla*<sub>CTXM</sub>-2, *bla*<sub>NDM</sub>, *aadA1*, *aadA2*, *sul1* | 0 | n/a | n/a |
| Raoultella sp. | 3 | none | 1 | *dhfr*-A15 | *bla*<sub>NDM</sub>, *bla*<sub>VIM</sub>, *aadA2*, *aacA7*, *sul1* | 0 | n/a | n/a |
| Enterobacter sp. | 2 | *bla*<sub>TEM</sub>, *bla*<sub>SHV</sub> | 1 | none | none | 0 | n/a | n/a |
| Pseudomonas sp. | 6 | *bla*<sub>TEM</sub>, *bla*<sub>CTX-M</sub> | 1 | *dhfr*-B3 | *bla*<sub>OXA</sub>-2, *bla*<sub>VIM</sub>-4, *bla*<sub>NDM</sub>-1, *bla*<sub>CMY</sub>, *aadA6*, *aacA8*, *aacA7*, *sul1* | 0 | n/a | n/a |
| Morganella sp. | 1 | *bla*<sub>TEM</sub> | 0 | n/a | n/a | 0 | n/a | n/a |
| Brenneria sp. | 4 | *bla*<sub>TEM</sub>, *bla*<sub>CTX-M</sub>, *bla*<sub>SHV</sub> | 0 | n/a | n/a | 0 | n/a | n/a |
| Comamonas sp. | 0 | n/a | 1 | none | none | 0 | n/a | n/a |
| Kluyvera sp. | 0 | n/a | 1 | *dhfr*-A21 | *bla*<sub>KPC</sub> | 0 | n/a | n/a |
| Fusobacterium sp. | 0 | n/a | 0 | n/a | n/a | 1 | *aadA5* | n/a |

### Characterization of bacterial communities by 16S rRNA sequencing

Of the 696 resistant bacterial isolates, 79 (11%) were unable to be speciated by 16s ribosomal RNA sequencing. From the remaining 617 resistant GNB isolates recovered from 102 samples, 11 unique genera were identified; AMP-resistant bacteria consisted of 6 distinct genera, TMP-SMX-resistant bacteria consisted of 7 distinct genera, GENT-resistant bacteria consisted of 2 genera, and COL-resistant bacteria consisted of 7 genera. The relative abundance of each bacterial genus is shown in Table 2.

Based on Shannon diversity index, COL-resistant colonies had the highest species diversity (H=1.9), followed by those resistant to AMP (H=1.2), TMP-SMX (H=0.8), and GENT (H=0.3) (Figure 2).

**Figure 2:**
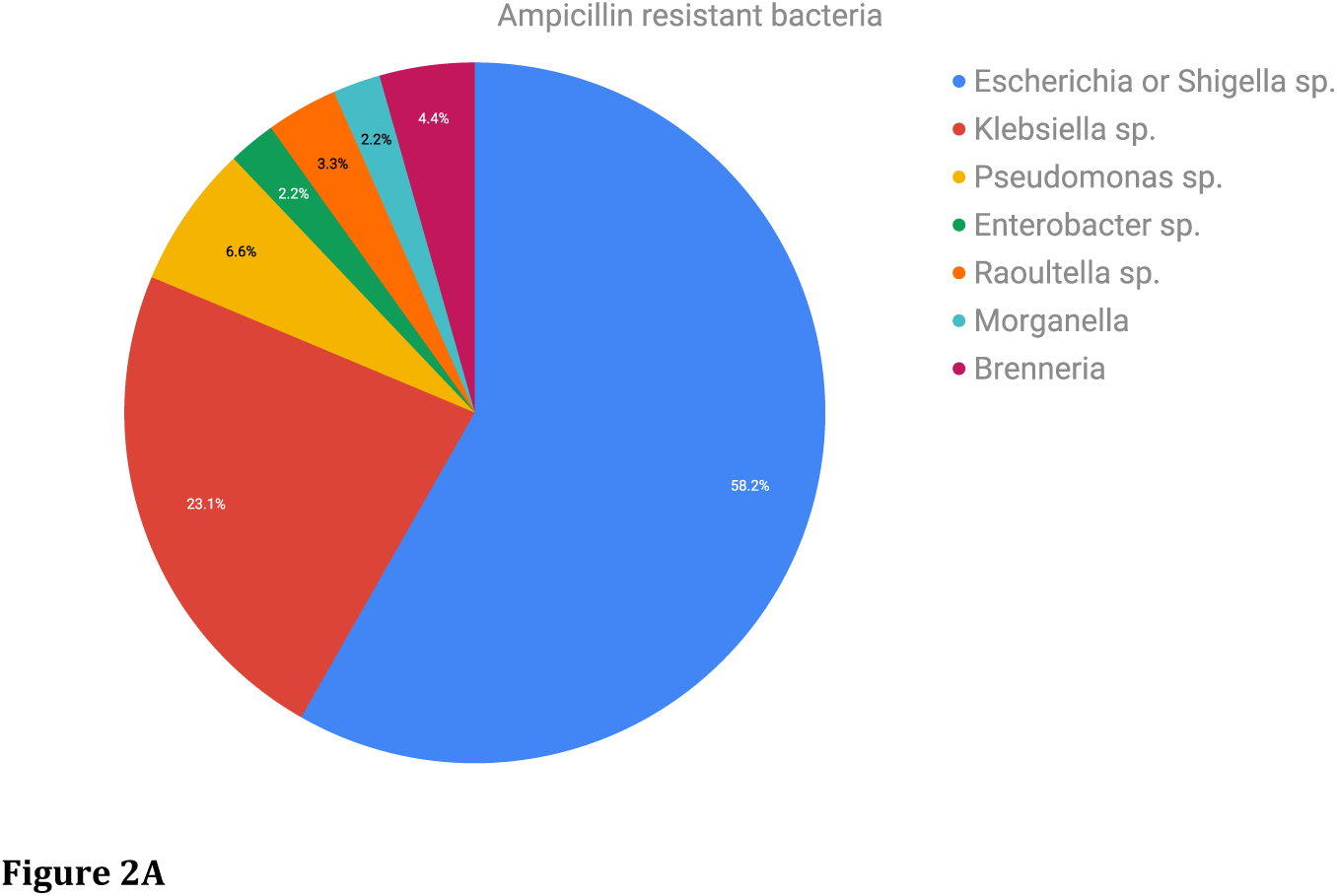
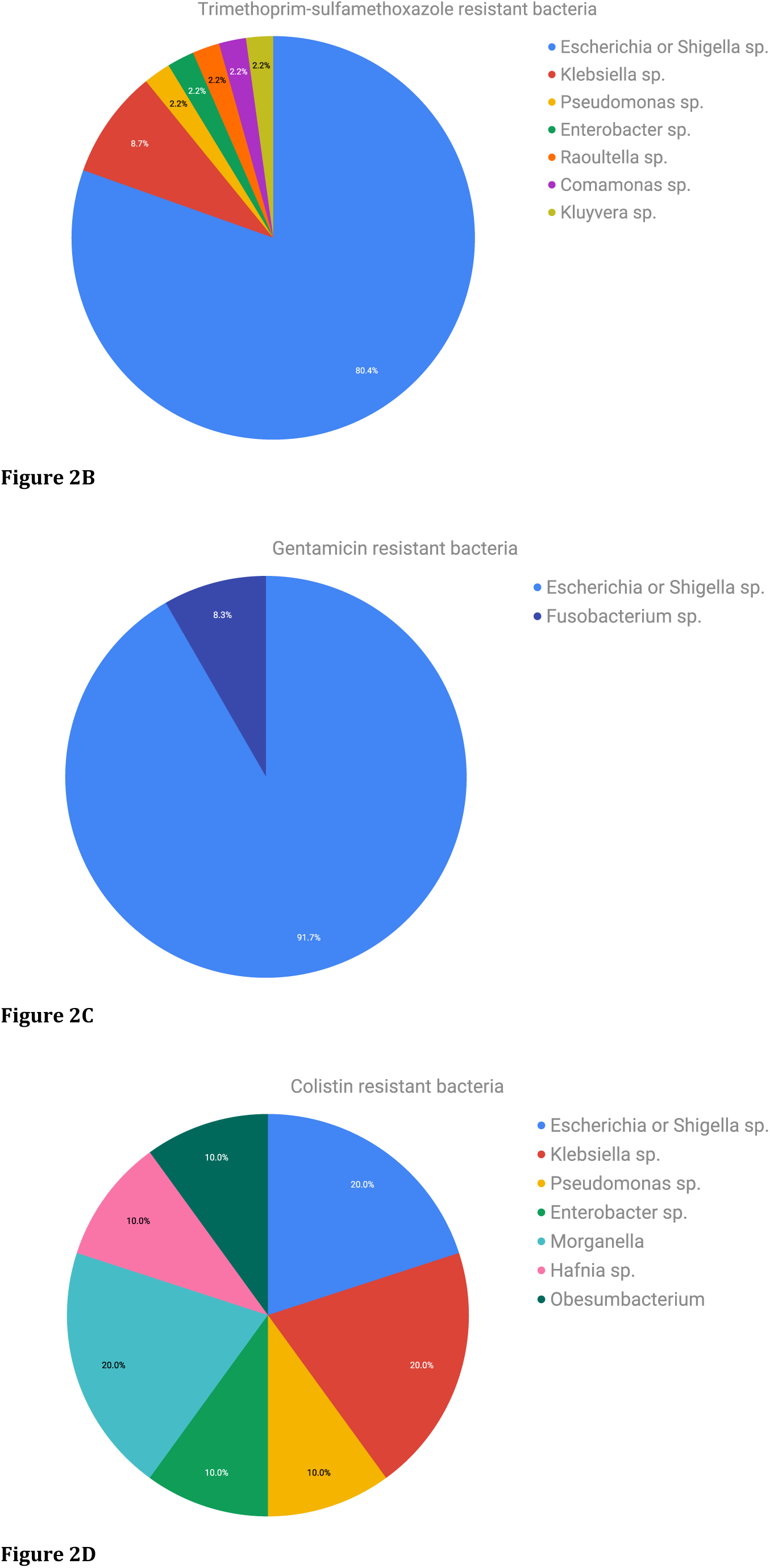
Relative frequency of genera represented by bacterial colonies exhibiting resistance to A) ampicillin, B) trimethoprim-sulfamethoxazole, C) gentamicin, and D) colistin.

Species identified in these samples included: *Escherichia spp (E*. *fergusonii*, *E. coli*, *Shigella flexneri*, *Shigella sonnei*, *Shigella dysenteriae*, *Shigella boydii)*, *Klebsiella spp (K. pneumoniae* or *quasipneumoniae*, *K. oxytoca*, *K. aerogenes*, *K. variicola)*, *Raoultella ornithinolytica* or *planticolla*, *Enterobacter spp* (*E. ludwigii*, *E. bugandensis*, *E. xiangfangensis*, *E. aerogenes*), *Pseudomonas aeruginosa*, *Morganella morganelli*, *Comamonas jiangduensis* or *terrigena*, *Kluyvera cryocrescens*, *Fusobacterium varium*, *Hafnia paralvei*, and *Obesumbacterium proteus*.

Although *Escherichia* sp. and *Klebsiella* sp. represented the majority of our fecal samples (79%), there were 21 samples that contained other types of bacteria (i.e. *Enterobacter* sp., *Pseudomonas* sp., *Shigella* sp., *Morganella* sp., *Raoultella* sp., and *Kluyvera* sp.). Among these 21 samples, 13 (62%) harbored the antibiotic resistance genes described above.

## Discussion

Our results demonstrate that there is an abundance of ARG (β-lactamase, *dhfr*, *aad*) carried by commensal GNB in healthy human intestine. Only 18% of rectal swab samples contained GNB that exhibited no phenotypic resistance to any of the four classes of drugs used to treat extraintestinal infections caused by enteric bacterial pathogens.

A previous study conducted at the same college community between 2016-2017 found that among 233 *E.coli* isolates cultured from human UTI cases, 97 (42%) were resistant to AMP, and 76 (78%) of these contained at least one β-lactamase gene. β-lactamase gene types from these isolates included *bla*_*TEM*_, *bla*_*CTX-M*_ group1, *bla*_*CTX-M*_ group9, *bla*_*SHV*_, and *bla*_*OXA*_ (43) (Table 3). Forty (17%) of the 233 *E. coli* isolates were resistant to TMP-SMX (43).

**Table 3:** Comparison of ARGs found in fecal samples (current study) compared to UTI samples obtained from the same population in 2016-2017 (Yamaji, 2018) and 2009 (Ajiboye, 2009). *P-value > .05 indicates a significant difference in genetic component between fecal samples and UPEC isolates.

| **Genes found in current study vs. clinical UPEC samples in same study population (Yamaji, 2018 and Ajiboye, 2009)** |  |  |  |
| --- | --- | --- | --- |
| **ARG** | **No. (%) fecal isolates (current study)** | **UPEC (Yamaji, 2018; Ajiboye, 2009)** | **P value** |
| *bla*TEM | 57 (74) | 72 (74) | 1.000 |
| *bla*CTX-M | 11 (14) | 6 (6) | 0.1209 |
| group 1 | 6 (8) | 4 (4) | 0.3409 |
| group 2 | 2 (3) | 0 (0) | 0.1944 |
| group 8/25 | 0 (0) | 0 (0) | 1.000 |
| group 9 | 3 (4) | 2 (2) | 0.656 |
| *bla*SHV | 19 (25) | 1 (1) | .0001* |
| *bla*OXA | 1 (1) | 7 (7) | 0.0784 |
| *dhfr*-A17 | 10 (21) | 38 (56) | .0005* |
| *dhfr*-A7 | 5 (11) | 3 (4) | 0.2621 |
| *dhfr*-A5 | 1 (2) | 3 (4) | 0.6438 |
| *dhfr*-A21 | 1 (2) | 0 (0) | 0.3982 |
| *dhfr*-A1 | 2 (4) | 2 (3) | 1.000 |
| *dhfr*-A15 | 1 (2) | 2 (3) | 1.000 |
| *dhfr*-B3 | 1 (2) | 0 (0) | 0.3982 |
| *dhfr*-A12 | 0 (0) | 0 (0) | 1.000 |
| *aad*A1 | 1 (8) | 17 (25) | 0.2783 |
| *aad*A5 | 5 (38) | 38 (56) | 0.3642 |
| *aad*A2 | 0 (0) | 5 (7) | 0.5868 |
| *aad*B | 1 (8) | 1 (1) | 0.2969 |

Another study of gene cassettes in human UTI *E*. *coli* isolates collected from the same college community from 1999-2001 found similar ARG harbored within the gene cassettes of these uropathogenic *E. coli*. *dhfr-A17* and *aadA5* were the most common cassette genes among these UTI isolates, which are the same ARG found in GNB in fecal samples in the current study. The findings of this study are summarized in Table 3.

A 2013 study by Adams-Sapper et al. found that *E. coli* isolates from patients with blood stream infection (BSI) at a San Francisco hospital contained β-lactamase genes similar to those of fecal GNB in our healthy volunteers (9); these included *bla*_CTX-M_ (CTX-M-15, CTX-M-14), *bla*_TEM_, and *bla*_OXA_. There was no significant difference between number of isolates containing CTX-M variants, or resistance to TMP-SMX in clinical BSI isolates compared to those in commensal bacteria in the present study (9). These observations demonstrate the potential for acquisition by extraintestinal pathogenic *E. coli* (ExPEC) of ARG via HGT from intestinal reservoirs.

Recent literature suggests geographic differences in gut ARG distribution. A 2008 study of fecal carriage of AMR in *E. coli* isolated from the gut of healthy adults in Paris found that only 2 (0.6%) of 332 fecal *E.coli* isolates contained ESBL CTX-M-15 gene (13). The present study identified at least 8 Escherichia spp isolates containing CTX-M-15, and 9 (9%) of 103 samples contained bacteria harboring CTX-M variants. These findings indicate a varying geographical distribution of the prevalence of AMR genes in healthy individuals, or a change in the prevalence of ARG over time, or both, which may be affected by differences or change in ingested food products containing AMR bacteria. Indeed, a 2018 study analyzed the resistome of 1,267 human gut samples from various countries. The findings indicated country-specific gut microbial signatures and significant differences in the gut resistome among different nationalities (44, 45). These studies, however, did not examine which gut GNB species carried the ARG.

In the present study, isolates from a given fecal sample usually contained the same set of ARG, even when the sample contained multiple species of bacteria. Forty-one (62%) of 76 AMP-resistant samples contained isolates in which all isolates of a given sample had the same composition of β-lactamase genes. These isolates belong to 8 different GNB species (*E. fergusonii*, *E. coli*, *E. marmotae*, *Shigella flexneri*, *Shigella sonnei*, *K. oxytoca*, *K. pneumoniae*, *K.dysenteriae)*. Our findings also suggest that *Klebsiella* species were more likely than non-*Klebsiella* species to possess SHV variants (P < .001), while *Escherichia* species were more likely than non-*Escherichia* species to harbor TEM variants (P<.001). This observation suggests high frequency of HGT of ARG among a host’s intestinal GNB microbiota when ARG enter the intestine.

Within integrons in both TMP/SMX- and GENT-resistant strains, all isolates within a given fecal sample contained the same ARG. This further supports the claim that genes are readily horizontally transferred to one another within the human gut, with integrons playing a particularly important role in this process (20). Gene cassettes present in the commensal bacteria analyzed in this study possessed sequences that were annotated to belong to carbapenemase and metallo-carbapenemase genes (*bla_KPC_, bla*_*NDM*_, *bla*_*IMP*_, *bla*_*VIM*_) as well as *mcr*-variant genes (Table 2). Interestingly, strains carrying cassette sequences annotated as *mcr*-variant were not phenotypically resistant to COL. Similarly, not all strains carrying cassette sequences annotated as beta-lactamase genes were resistant to AMP.

In the study of *E. coli* isolates by Ajiboye et al., pathogenic strains from food animal origins and human UTI isolates shared many of the same gene cassette sequences (19), as well as those found in commensal bacteria in the current study (Table 3). Such mobile elements carrying ARG may be introduced into the human intestine by food contaminated with pathogenic as well commensal bacteria harboring these ARG, creating an abundant intestinal reservoir for ARG that then may get transferred into GNB strains that cause extraintestinal infections.

The scope of this study was limited to analysis of aerobic or microaerophilic GNB species. This design was intentional as most mobile ARG found in GNB pathogens are restricted to these species. However, it must be noted that the majority of gut microbial organisms are anaerobic, and thus the findings of our study may underestimate the frequency of HGT of ARG that may originate in anaerobes. In addition, our analysis of integron sequences was limited to class 1 integrons and to strains resistant to GENT or TMP-SMX, which would underestimate the prevalence of ARGs in integrons in fecal samples from this healthy college study population. Many of the ARG in this study’s AMP-resistant strains may be harbored by integrons, together with other ARG cassettes. Further studies that investigate HGT between a broader range of microbial species must be conducted to wholly reveal the reservoir of ARG among intestinal bacterial organisms, and the implications for human health.

This study also utilized a screening method to identify ARG based on the patterns of phenotypic resistance. It is possible that the GNB isolates harbored ARG that conferred resistance to antimicrobial agents we did not use for screening. Further investigations into the resistome separated by GNB species in commensal bacteria will illuminate the implications of HGT between bacterial species.

Our study showed abundance and diversity of resistance genes among healthy adults in a particular college community. While these results may not be generalizable to other communities, it highlights the need for community-based comparisons between commensal and pathogenic bacteria. These observations made in a non-healthcare environment raise questions about how these ARG were introduced into the intestines of people residing in this community, including food and environment as sources. It calls for further investigation into possible risk factors for acquisition of ARG other than just exposures to antibiotics, as they clearly pose a risk to public health. The findings in this study illuminate the relationship between gut microbial ARG content and infections such as UTI caused by Gram-negative pathogens. These observations may have global implications for the spread of antibiotic resistance through mobile genetic elements, as infections become increasingly difficult to treat with current antibiotic therapies.

## Acknowledgements

We would like to thank Reina Yamaji for her assistance in the study design process, as well as Clarissa Araujo Bourges for her consistent support in classification of study isolates. We would also like to thank the University instructors who were instrumental in recruitment for this study. We would also like to the University Department of Environmental Health and Safety for their assistance in ensuring safe shipment of biological specimens.

## Financial support

This study was supported by Centers for Disease Control and Prevention program to combat antibiotic resistance under BAA number 200-2016-91939.

## Conflicts of interest

All authors: no conflicts.

## Supplemental Material

**Table S1:**
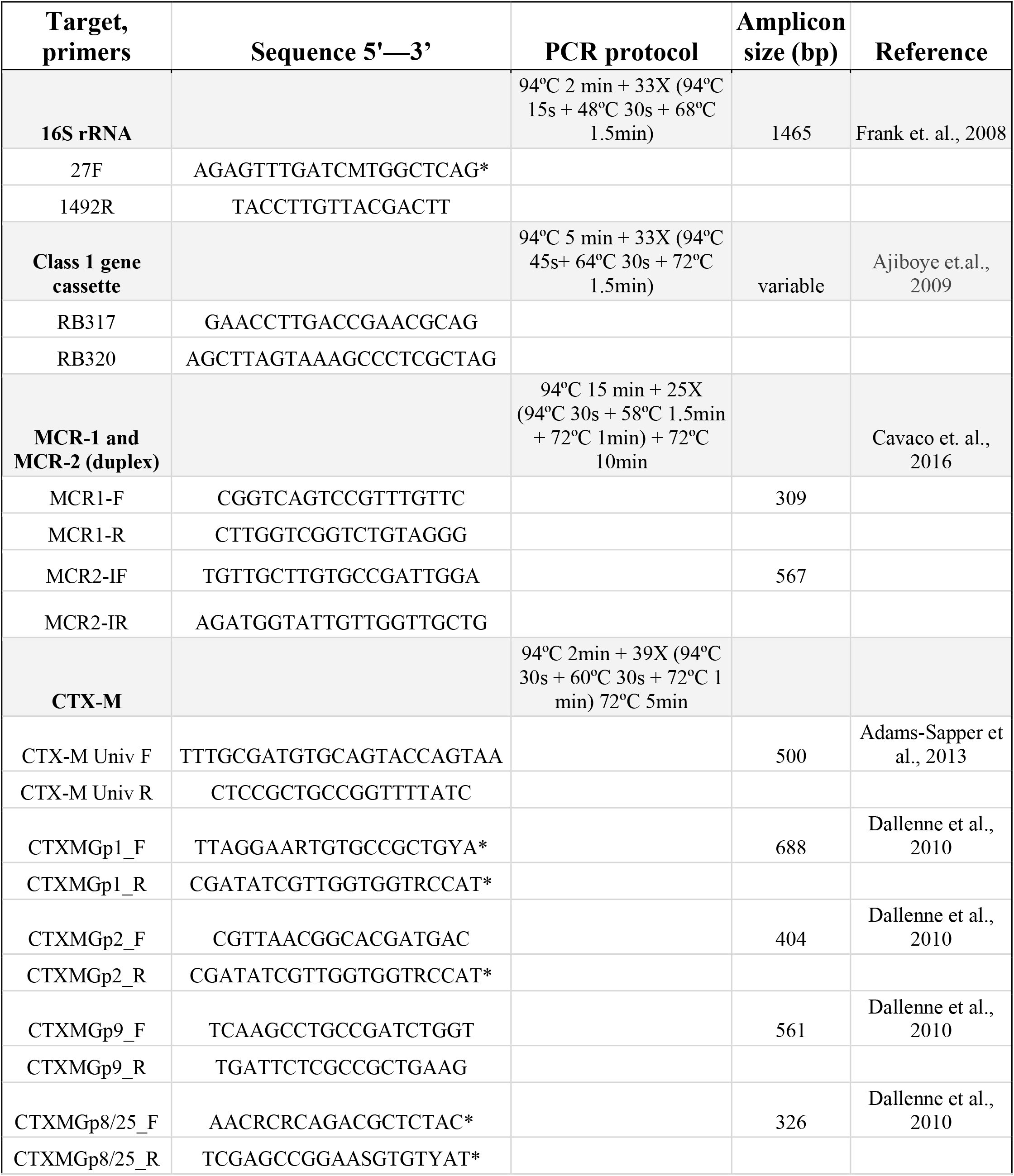
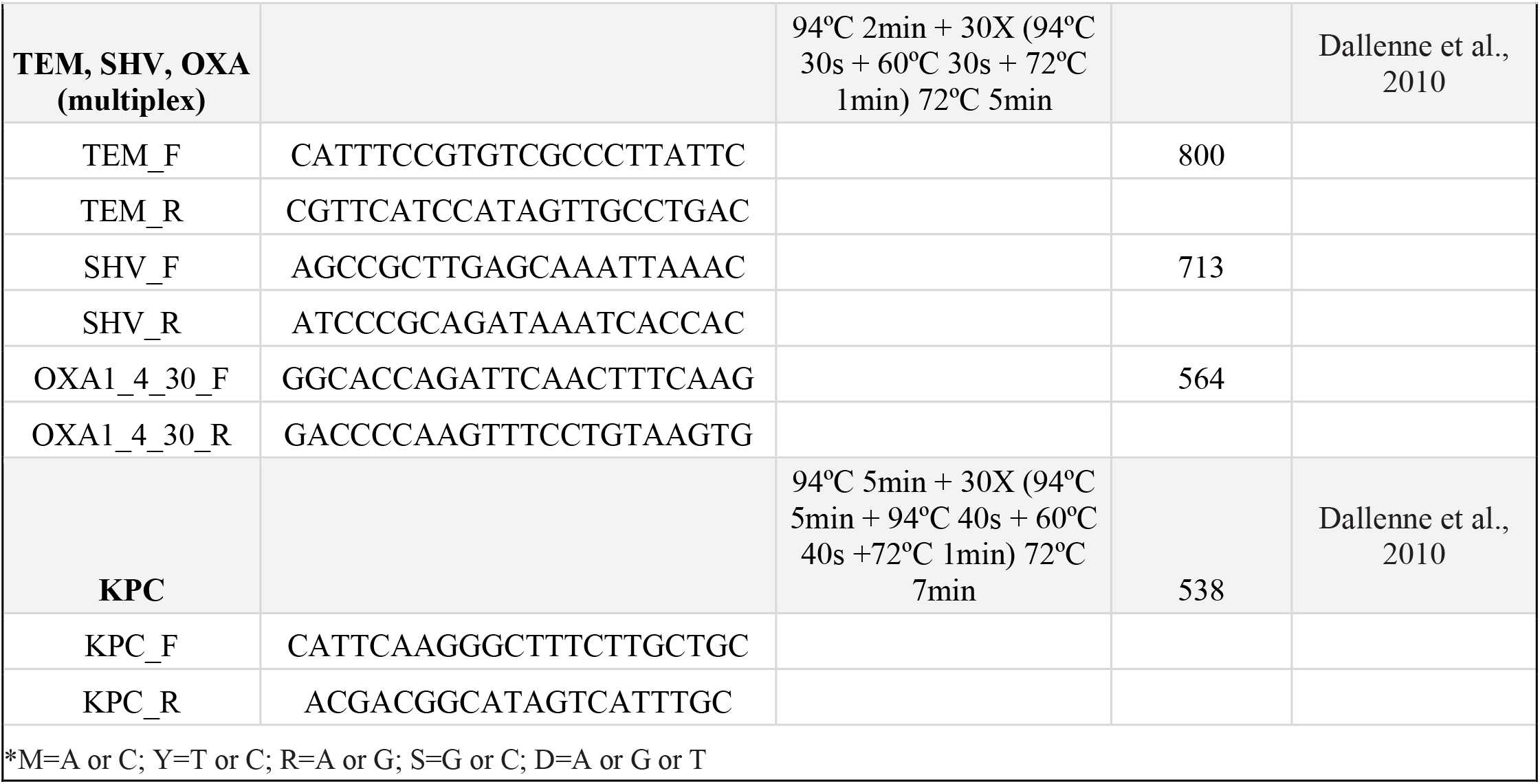
Polymerase chain reaction (PCR) primers and conditions.

**Table S2:**
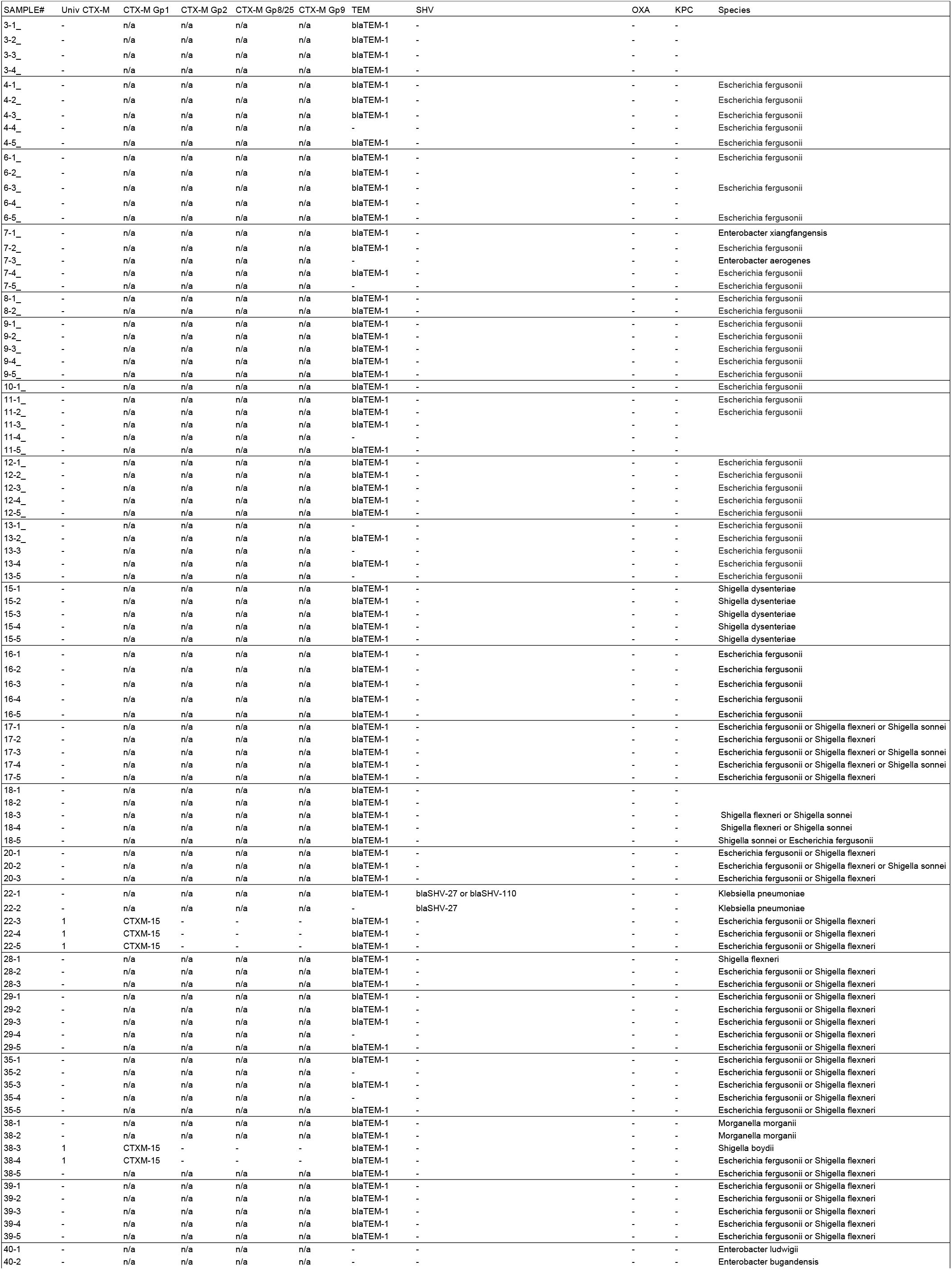
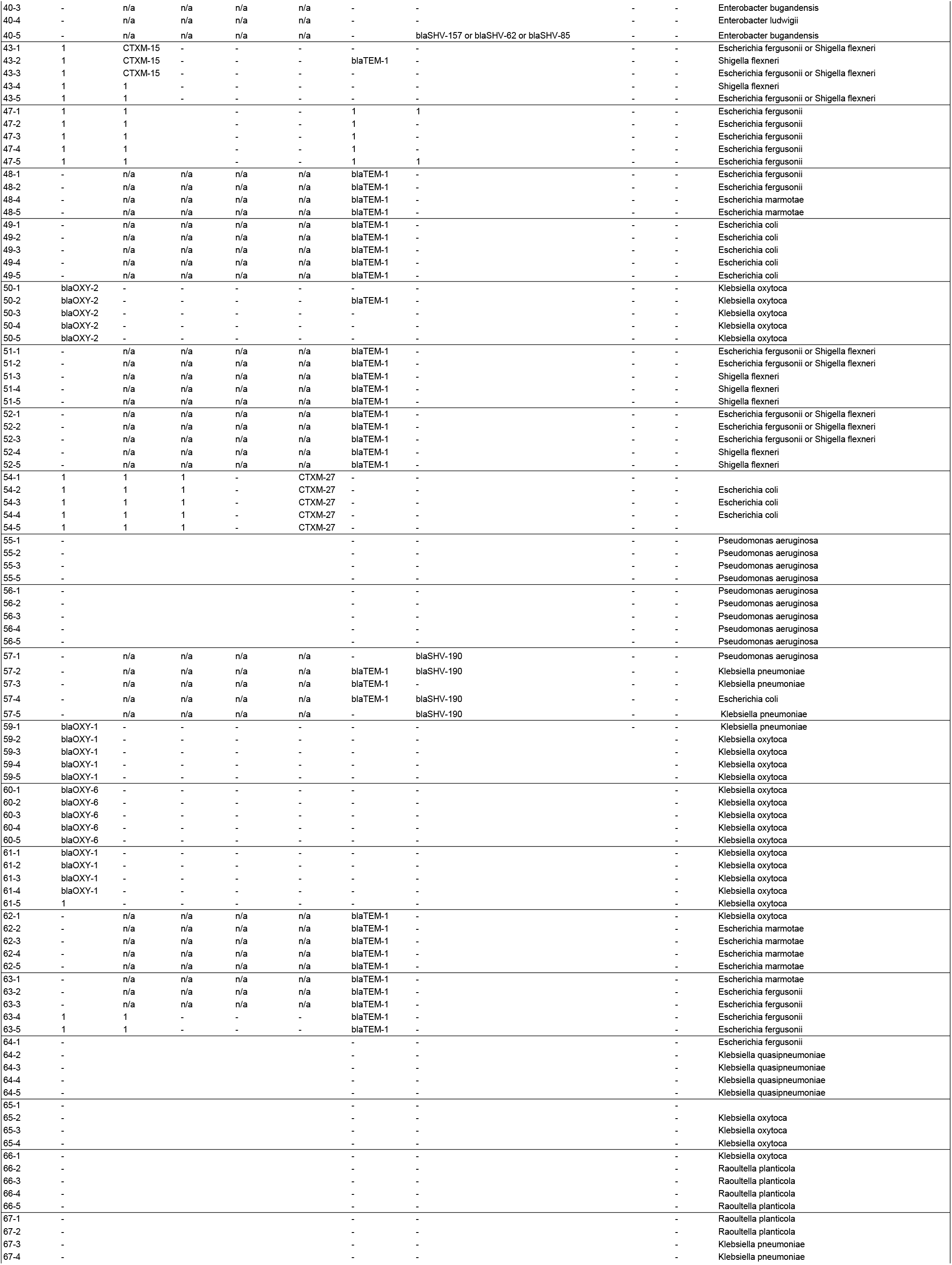
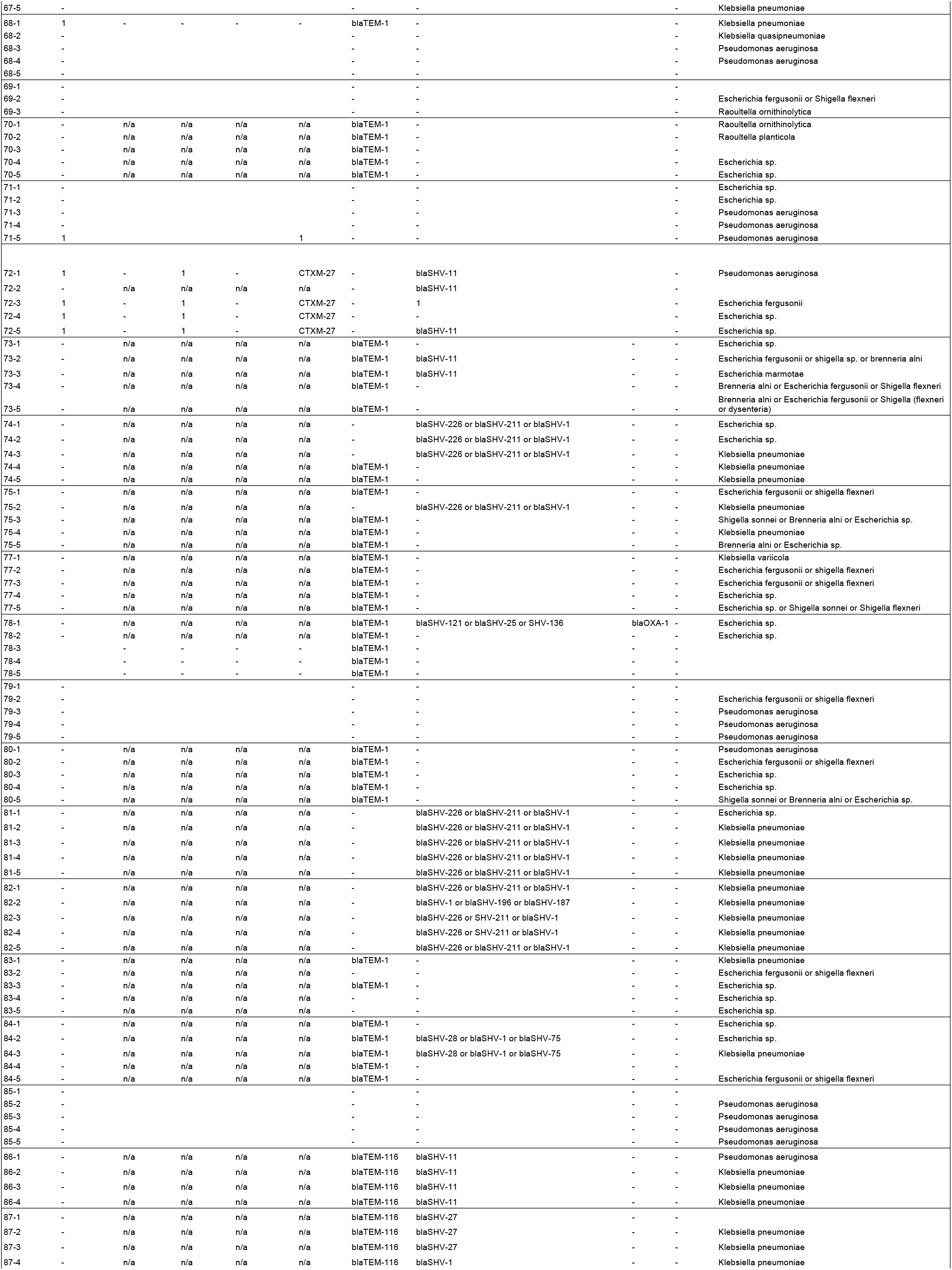
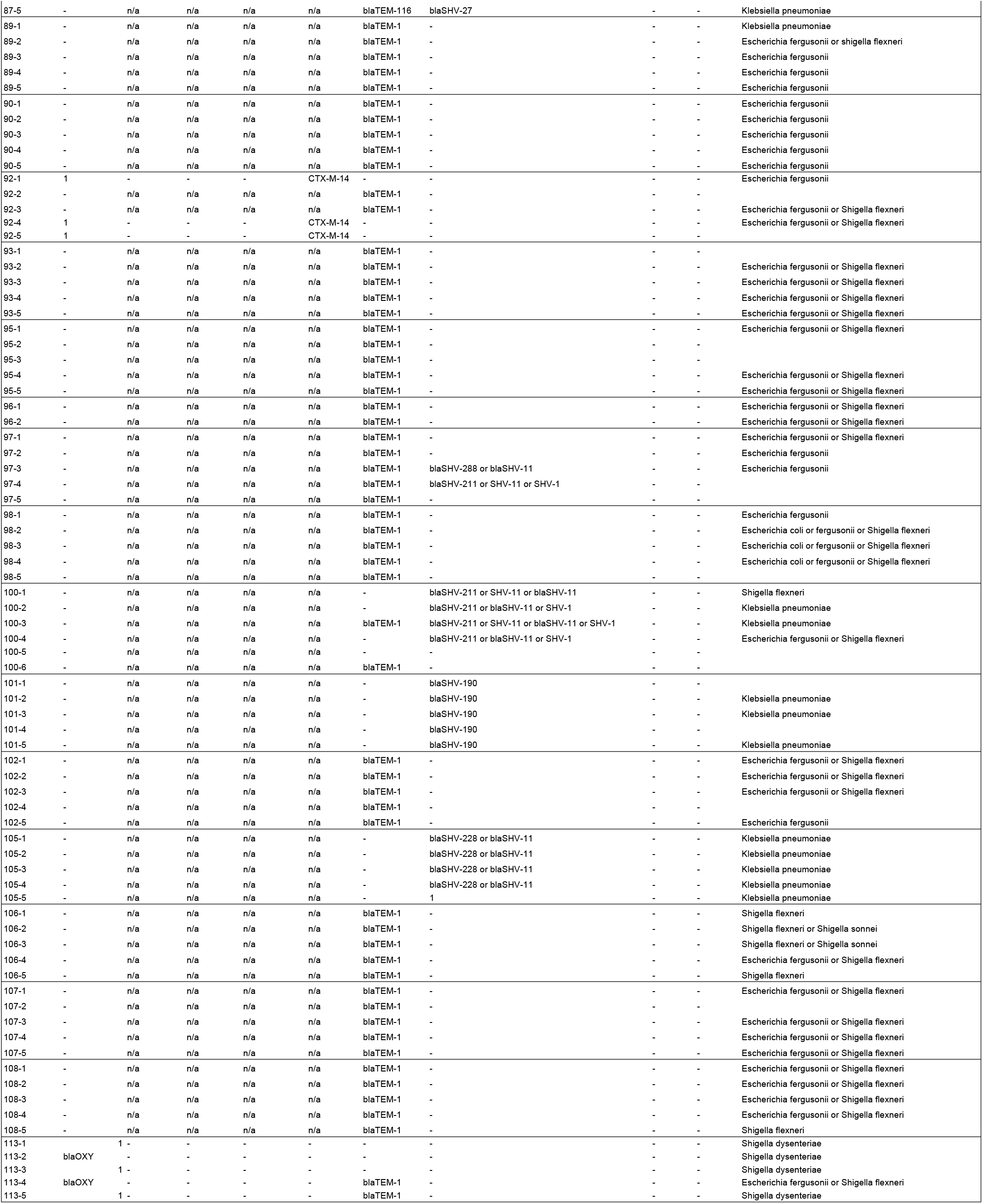
ARG and species associated with AMP-resistant colonies.

**Table S3:**
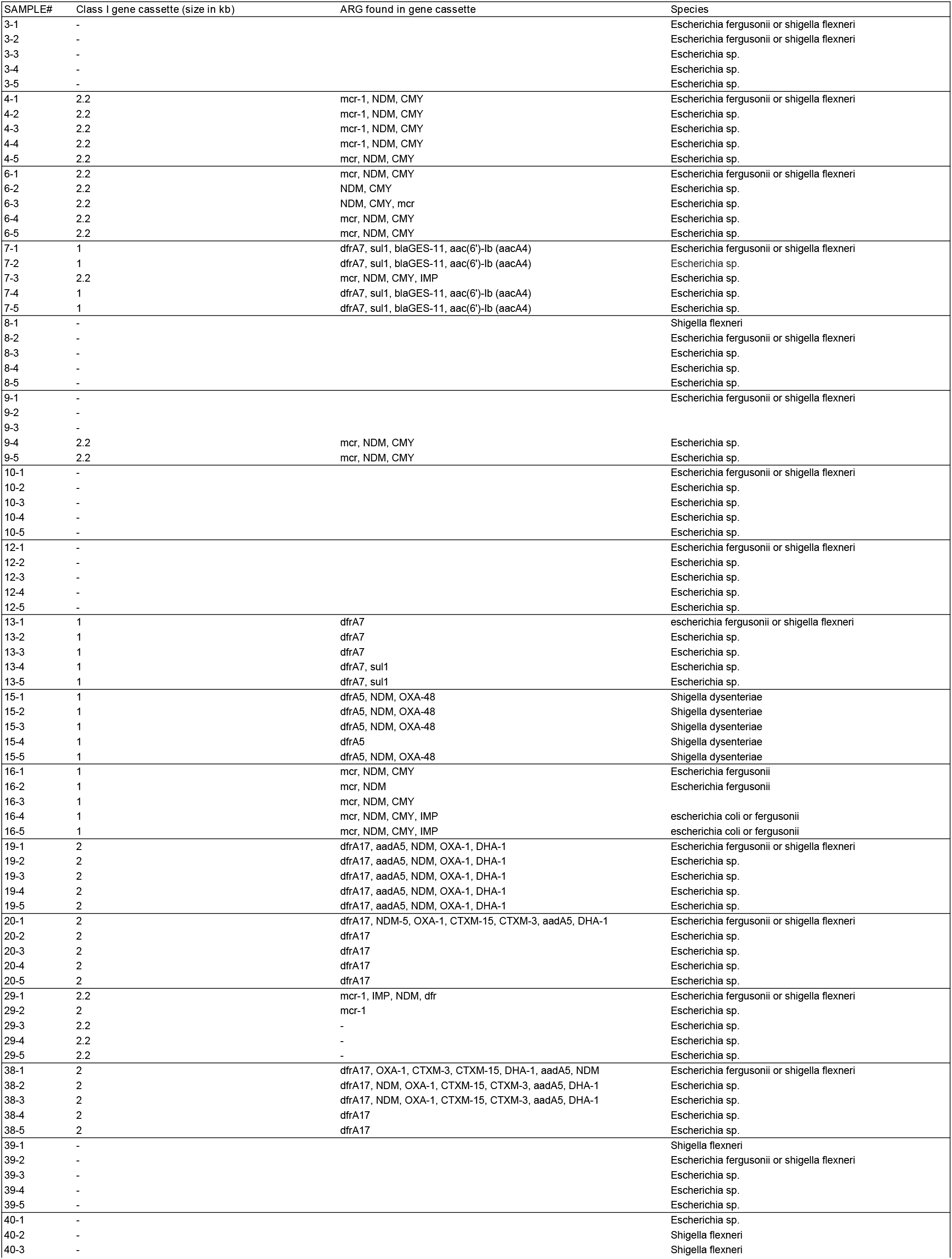
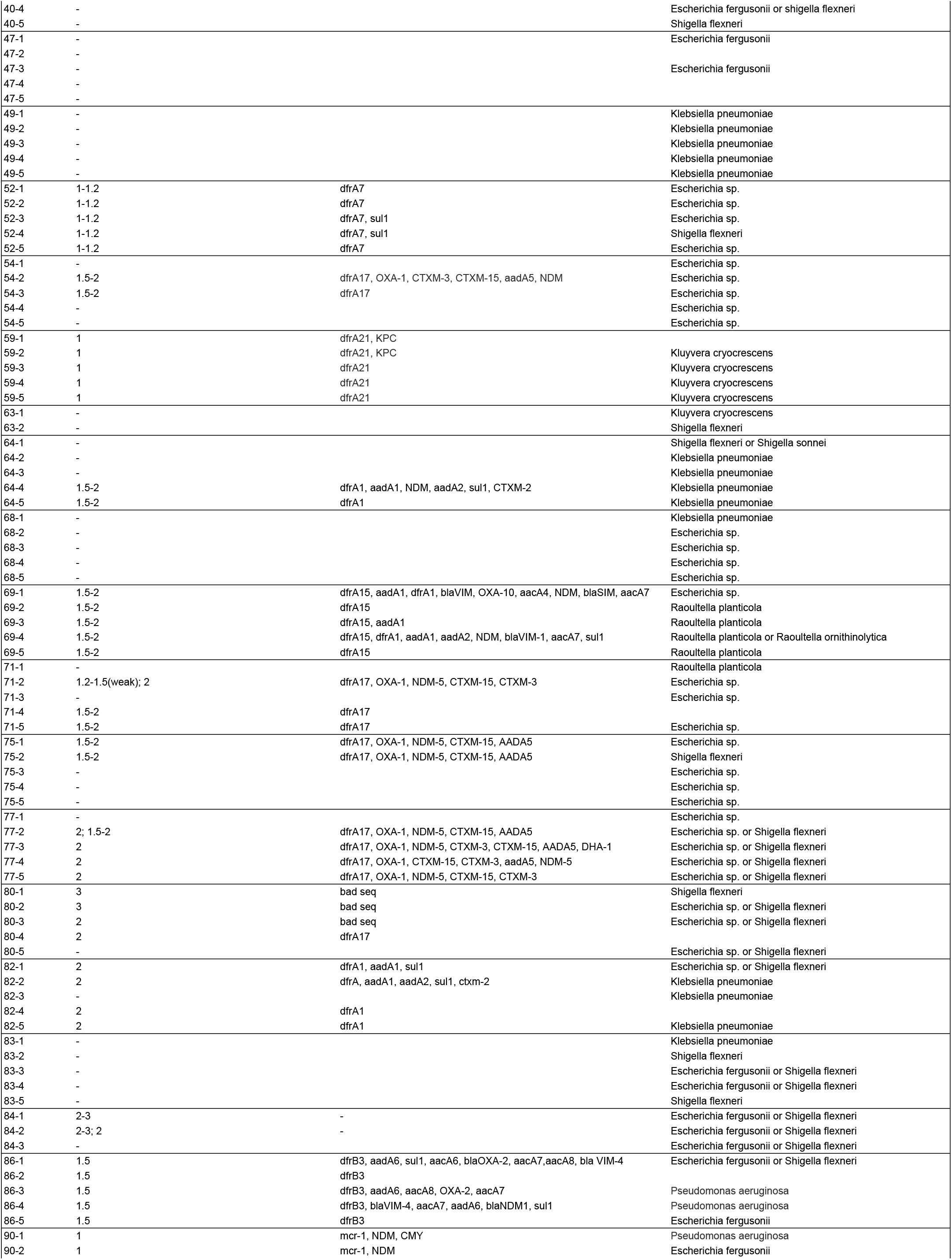
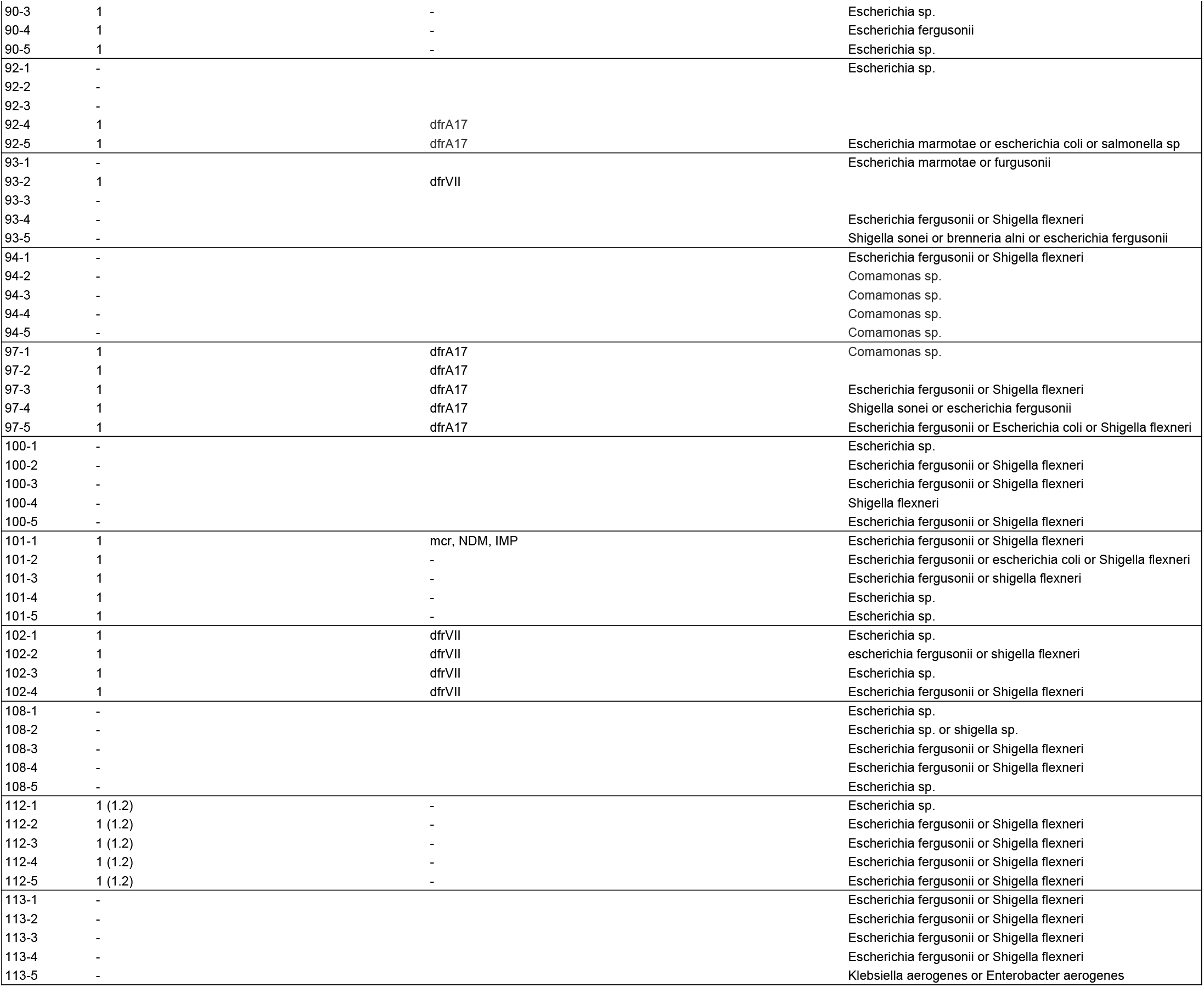
ARG and species associated with TMP/SMX-resistant colonies.

**Table S4:** ARG and species associated with GENT-resistant colonies.

| SAMPLE# | Class I gene cassette | ARG found in gene cassette | Species |
| --- | --- | --- | --- |
| 19-1 | 1 | aadA5, NDM-5, OXA-1, CTXM-15 | Escherichia fergusonii or shigella flexneri |
| 19-2 | 1 | aadA5, OXA-1, CTXM-15, CTXM-3, NDM-9, NDM-5 | Escherichia fergusonii or shigella flexneri |
| 20-1 | 1 | aadA5, OXA-1, CTXM-15, CTXM-3, NDM-9, NDM-5 | Escherichia fergusonii or shigella flexneri |
| 20-2 | 1 | aadA5, OXA-1, CTXM-15, CTXM-3, NDM-9, NDM-5 |  |
| 20-3 | 1 | aadA5, OXA-1, CTXM-15, CTXM-3, NDM-9, NDM-5 | Escherichia fergusonii or shigella flexneri |
| 20-4 | 1 | aadA5, OXA-1, CTXM-15, CTXM-3, NDM-9, NDM-5 | Escherichia fergusonii or shigella flexneri |
| 20-5 | 1 | aadA5, OXA-1, CTXM-15, CTXM-3, NDM-9, NDM-5 | Escherichia fergusonii or shigella flexneri |
| 38-1 | 1 | aadA5, OXA-1, CTXM-15, CTXM-3, NDM-9, NDM-5 | Escherichia fergusonii or shigella flexneri |
| 38-2 | 1 | aadA5, OXA-1, CTXM-15, CTXM-3, NDM-9, NDM-5 | Escherichia fergusonii or shigella flexneri |
| 38-3 | 1 | aadA5, OXA-1, CTXM-15, CTXM-3, NDM-9, NDM-5 | Escherichia fergusonii or shigella flexneri |
| 38-4 | 1 | aadA5, OXA-1, CTXM-15, CTXM-3, NDM-9, NDM-5 | Escherichia fergusonii or shigella flexneri |
| 38-5 | 1 | aadA5, OXA-1, CTXM-15, CTXM-3, NDM-9, NDM-5 | Escherichia fergusonii or shigella flexneri |
| 47-1 | - |  | escherichia fergusonii |
| 47-2 | - |  | escherichia fergusonii |
| 47-3 | - |  | escherichia fergusonii |
| 47-4 | - |  | escherichia fergusonii |
| 47-5 | - |  | escherichia fergusonii |
| 57-1 | - |  | escherichia fergusonii |
| 57-2 | - |  | escherichia fergusonii |
| 57-3 | - |  | escherichia fergusonii |
| 57-4 | - |  | escherichia fergusonii |
| 57-5 | - |  | escherichia fergusonii |
| 68-1 | - |  | Escherichia coli |
| 68-2 | - |  | Escherichia coli |
| 68-3 | - |  | Escherichia coli |
| 68-4 | - |  | Escherichia fergusonii or shigella flexneri |
| 68-5 | - |  | Escherichia sp. or Shigella sonnei or shigella flexneri |
| 70-1 | 1 | aadA5 | Shigella flexneri |
| 70-2 | 1 | aadA5 | Shigella flexneri |
| 70-4 | 1 | aadA5 | Shigella flexneri |
| 70-5 | 1 | aadA5 | Shigella flexneri |
| 73-1 | 1 | NDM, IMP, MCR, CMY | Escherichia fergusonii or shigella flexneri |
| 73-2 | 1 | NDM, IMP | Escherichia marmotae |
| 73-3 | 1 | NDM, IMP, CMY | Escherichia fergusonii or shigella flexneri or Escherichia marmotae |
| 73-4 | 1 | AADA2, NDM, CMY, IMP | Escherichia marmotae |
| 73-5 | 1 | NDM, IMP, CMY | Escherichia fergusonii or shigella flexneri or Escherichia marmotae |
| 74-1 | - |  | Shigella sonnei or Escherichia fergusonii or shigella flexneri |
| 74-2 | - |  | Shigella sonnei or Escherichia fergusonii or shigella flexneri |
| 74-3 | - |  | Shigella sonnei or Escherichia fergusonii or shigella flexneri |
| 74-4 | - |  | Shigella sonnei or Escherichia fergusonii or shigella flexneri |
| 74-5 | - |  | Escherichia fergusonii or shigella flexneri |
| 75-1 | 1 | aadB, aadA1 | Escherichia fergusonii or shigella flexneri or shigella sonnei or brenneria alni |
| 75-2 | 1 | aadB, aadA1 | Shigella flexneri |
| 75-3 | 1 | aadB, aadA1 | shigella flexneri |
| 75-4 | 1 | aadB, aadA1 | Escherichia fergusonii or shigella flexneri |
| 75-5 | 1 | aadB, aadA1 | Escherichia fergusonii or shigella flexneri |
| 86-1 | - |  | Escherichia fergusonii or shigella flexneri |
| 86-2 | - |  | Escherichia fergusonii or shigella flexneri |
| 86-3 | - |  | Escherichia fergusonii or shigella flexneri |
| 86-4 | - |  | Escherichia fergusonii or shigella flexneri |
| 86-5 | - |  | Escherichia fergusonii or shigella flexneri |
| 106-1 | 1 | aadA5 | Fusobacterium varium |
| 106-2 | 1 | aadA5 |  |
| 106-3 | 1 | aadA5 | Escherichia fergusonii or shigella flexneri or shigella dysenteriae |
| 113-1 | - |  |  |
| 113-2 | - |  |  |
| 113-3 | - |  |  |
| 113-4 | - |  |  |
| 113-5 | - |  |  |

**Table S5:** ARG and species associated with COL-resistant colonies.

| **SAMPLE#** | **mcr-1** | **mcr-2** | **Species** |
| --- | --- | --- | --- |
| 16-1 | - | - | *Escherichia fergusonii* |
| 17-1 | - | - | *Enterobacter* sp. |
| 18-1 | - | - | *Escherichia* sp. |
| 18-2 | - | - | *Escherichia* sp. |
| 26-1 | - | - | *Klebsiella pneumoniae* |
| 26-2 | - | - | *Klebsiella pneumoniae* |
| 26-3 | - | - | *Klebsiella pneumoniae* |
| 26-4 | - | - | *Klebsiella pneumoniae* |
| 26-5 | - | - | *Klebsiella pneumoniae* |
| 40-1 | - | - | *Hafnia paralvei* |
| 40-2 | - | - | *Hafnia paralvei* |
| 40-3 | - | - | *Hafnia paralvei* |
| 40-4 | - | - | *Hafnia paralvei* |
| 40-5 | - | - | *Hafnia paralvei* |
| 42-1 | - | - | *Escherichia coli* or *fergusonii* |
| 42-2 | - | - | *Escherichia coli* or *fergusonii* |
| 42-3 | - | - | *Escherichia coli* |
| 42-4 | - | - | *Escherichia fergusonii* or *coli* |
| 42-5 | - | - | *Escherichia coli* |
| 47-1 | - | - | *Klebsiella pneumoniae* |
| 47-2 | - | - | *Klebsiella pneumoniae* |
| 47-3 | - | - | *Klebsiella pneumoniae* |
| 47-4 | - | - |  |
| 47-5 | - | - | *Klebsiella pneumoniae* |
| 100-1 | - | - | *Morganella morganii* |
| 100-2 | - | - | *Morganella morganii* |
| 100-3 | - | - | *Morganella morganii* |
| 100-4 | - | - | *Morganella morganii* |
| 100-5 | - | - | *Pseudomonas aeruginosa* |
| 105-1 | - | - | *Obesumbacterium proteus* strain |
| 105-2 | - | - | *Obesumbacterium proteus* strain |
| 105-3 | - | - | *Obesumbacterium proteus* strain |
| 105-4 | - | - | *Obesumbacterium proteus* strain |
| 105-5 | - | - | *Obesumbacterium proteus* strain |
| 113-1 | - | - | *Morganella morganii* |
| 113-2 | - | - | *Morganella morganii* |
| 113-3 | - | - | *Morganella morganii* |
| 113-4 | - | - | *Morganella morganii* |
| 113-5 | - | - | *Morganella morganii* |

